# Overcoming the Limits of Traditional Rate Calculations from Sparse Concentration Data: A Probabilistic Framework for Bioprocess Modeling

**DOI:** 10.1101/2025.09.30.679468

**Authors:** Anne Richelle, David Andersson, Anton Vernersson, Olivier Cloarec, Johan Trygg

## Abstract

Accurate estimation of growth and metabolic rates is essential for understanding and optimizing bioprocesses, yet traditional methods often fail when faced with sparse or noisy concentration data. We present a probabilistic framework based on Bayesian inference and Nested Sampling that addresses these challenges by integrating biological knowledge directly into the model structure. The approach transforms raw concentration measurements into pseudo-concentrations that account for distortions caused by bioreactor volume changes (e.g., feed additions, sample withdrawals), and models metabolic rates as linear combinations of basis functions to yield continuous rate profiles from discrete data. Using in-silico simulations, we evaluated the framework under a range of experimental conditions and compared its performance with a conventional rate calculation method. We further analyzed the influence of key experimental design parameters - sampling frequency, sample volume, and measurement noise - on both rate estimation accuracy and concentration reconstruction quality. Results demonstrate that the proposed framework delivers accurate, robust metabolic rate estimates even under severe data sparsity and noise, offering a powerful tool for improving bioprocess characterization and optimization.

## 1. Introduction

In the context of bioprocesses, accurately estimating growth and metabolic rates is essential for understanding biological systems and optimizing production processes (Richelle et al., 2020). However, this task remains challenging due to the sparse and noisy nature of concentration data, which are typically obtained from discrete, low-frequency measurements using online analytical devices such as optical sensors and cell counters. These limitations are compounded by the inherent stochasticity of biological systems, which introduces additional variability and complexity (Sonnleitner et al., 2000; Zahel et al., 2016; Bayer et al., 2020).

Traditional approaches for rate estimation, such as stepwise integration between consecutive sampling time points, assume constant rates over each interval regardless of its length (Wechselberger et al., 2013; Bayer et al., 2020). While straightforward, this simplification often fails to capture dynamic transitions in nonlinear biological systems. Smoothing techniques such as moving average filters have been applied to mitigate fluctuations but tend to obscure genuine dynamics and may propagate errors (Paulsson et al., 2014). More advanced interpolation techniques - including polynomial, cubic splines, and radial basis functions - offer greater flexibility but require careful parameterization to avoid artefacts. In particular, spline-based methods such as cubic smoothing splines (Bayer et al., 2020) and B-splines applied in dynamic metabolic flux analysis (Martínez et al., 2015) have shown promise in generating continuous rate profiles, yet they remain sensitive to oversmoothing and depend on frequent, high-quality measurements to constrain the fit. Probabilistic approaches, such as Gaussian process regression, extend these ideas by quantifying uncertainty in derivative inference (Swain et al., 2016), but also rely on dense datasets and are less reliable under the sparse, noisy measurement regimes typical of fed-batch bioprocesses.

Despite these advances, existing methods remain limited when applied to the sparse and noisy datasets typical of bioprocess monitoring. Spline-based approaches require frequent, high-quality measurements to avoid artefacts, while probabilistic regressions such as Gaussian processes lose reliability under irregular sampling. There is therefore a need for frameworks that can robustly estimate biological rates under realistic data constraints, while explicitly propagating uncertainty from raw measurements to derived rates to ensure that predictions remain both informative and reliable.

To address these challenges, we propose a Bayesian inference framework that models metabolic consumption rates as a linear combination of basis functions. These basis functions are selected to reflect known growth stages of cell cultures, capturing the typical transition from lag to exponential and stationary phases. This approach transforms sparse concentration measurements into continuous rate profiles by incorporating biological insight directly into the structure of the model.

We implement Bayesian parameter inference using Nested Sampling, a powerful algorithm developed by Skilling (2004, 2006) that not only samples from the posterior distribution but also computes the Bayesian evidence, enabling formal model comparison. Compared to traditional Markov Chain Monte Carlo (MCMC) methods - such as Gibbs sampling or Hamiltonian Monte Carlo - Nested Sampling offers advantages in dealing with multimodal posteriors and exploring complex parameter spaces efficiently (Feroz et al., 2009; Speagle, 2020; Buchner, 2021).

Our framework further incorporates a pseudo-batch transformation to account for volume changes in concentration data, improving comparability across time points. The resulting method enables robust estimation of growth and metabolic rates from sparse and noisy datasets, while also supporting principled model selection using the Bayesian evidence. As computational demands grow and modeling complexity increases, this approach provides a scalable, data-efficient, and probabilistically grounded tool for improving the accuracy and interpretability of bioprocess models.

The remainder of this manuscript is structured as follows. Section 2 (Materials and Methods) details the proposed probabilistic framework, including its mathematical formulation and the integration of Bayesian inference with Nested Sampling. Section 3 (Results) presents in-silico simulation studies evaluating the framework’s ability to estimate growth and metabolic rates under a range of experimental conditions, alongside an analysis of how sampling frequency, sample volume, and measurement noise influence both rate estimation accuracy and concentration reconstruction. Section 4 (Discussion) interprets these findings in the context of bioprocess engineering, highlighting the framework’s potential for future integration into real-time control and short-horizon predictive modeling, and outlining methodological extensions to enhance its applicability in advanced process monitoring strategies.

## 2. Materials and Methods

The approach proposed creates models using linear combinations of differentiable basis functions and applies Bayesian inference to update knowledge of model parameters, with the goal of generating credible approximations of specific consumption. A general overview of the method can be seen in Figure 1, which illustrates how data is pre-processed, interpolated, and used to infer model parameters using the various components of the inference framework, referred to in this work as ‘MetRaC’ for ‘Metabolic Rate Calculator’.

**Fig 1.**
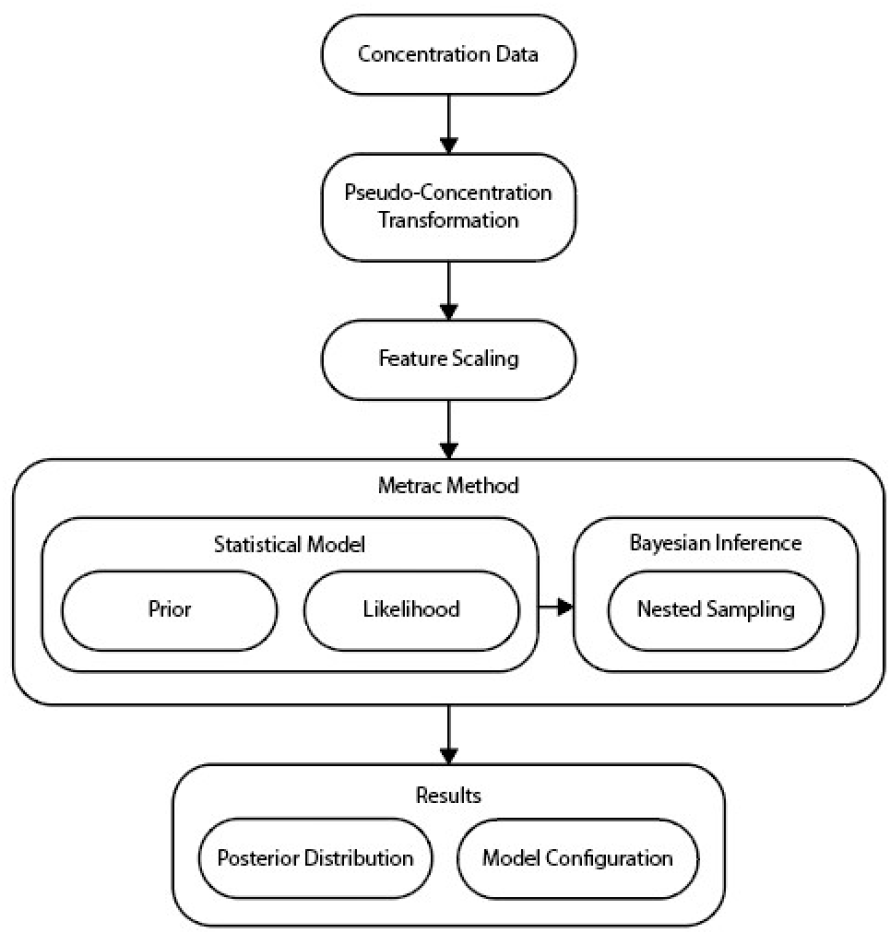
Overview of the framework components for estimating cell-specific metabolic rates.

### 2.1. Preprocessing

#### Pseudo-concentrations calculations

In the context of cell culture bioprocessing, accurately computing metabolic rates from concentration data requires careful consideration of discrete volume additions. These additions, which may contain varying concentrations of one or more components, can significantly impact all measurements within the working volume. To ensure precise assessment of the rates of change in component concentration, data from these volume addition events should be excluded. This exclusion also extends to any external processing events that alter the working volume (for example, bleeding feed during continuous culture operation).

For fed-batch processes, pseudo-counterparts of metabolite concentrations and viable cell density are derived using the method outlined in Hesselberg-Thomsen (2024), where the batchwise accumulated dilution factor (ADF), γ, is calculated as the product of relative changes in reactor volume.

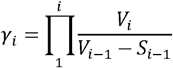

here, *i* denotes the time index *i* = 1,2,…,*N*_*t*_ (for *N*_*t*_ timesteps), *V* is the volume prior to sampling and *S* is the sample volume. If measurements are taken by a probe that does not withdraw a sample, then *S* = 0. Any action on the reactor that is not a volume-changing, concentration-preserving, and instantaneous operation will have its *S* = 0. The transformation from metabolite concentration *M* to pseudo-concentration *M*\* at a measurement time *i*, for a metabolite indexed by *i* = 1,2,…,*N*_*M*_ (for *N*_*M*_ number of metabolites), is given by:

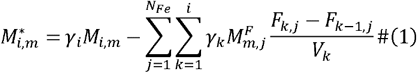

Here, feed additions are indexed by *i* = 1,2,…,*N*_*Fe*_ (for *N*_*Fe*_ number of feeds), *k* denotes the time index ranging from 1 to *i (*measurement times*), M*^*F*^ is the concentration of the metabolite in the feedstock, and *F* is the cumulated feed volume. [*F*_*k,j*_ − *F*_*k−*1,*j*_] is therefore the feed-induced difference in volume between the current time index and the preceding one (i.e., accumulated change due to feeding), which, depending on how often measurements are taken, could include volumes for more than 1 bolus feed (and could have occurred over a period where one or more samples are also taken). For viable cell density, *X*, there are no feed additions; hence, the transformation is given by:

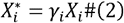

where *X*\* is the pseudo-density. In order to simplify notation, metabolites are not indexed in the following sections.

Figure 2 is an example of the pseudo-concentration transformation process, showing the transformation of glucose and lactate concentration in a fed-batch culture (upper panels) to a pseudo-concentration (lower panels) varying over time, using the in-silico dataset generated as presented in Section 2.7. Such a transformation allows for the exclusion of discrete volume additions from the data.

**Fig 2.**
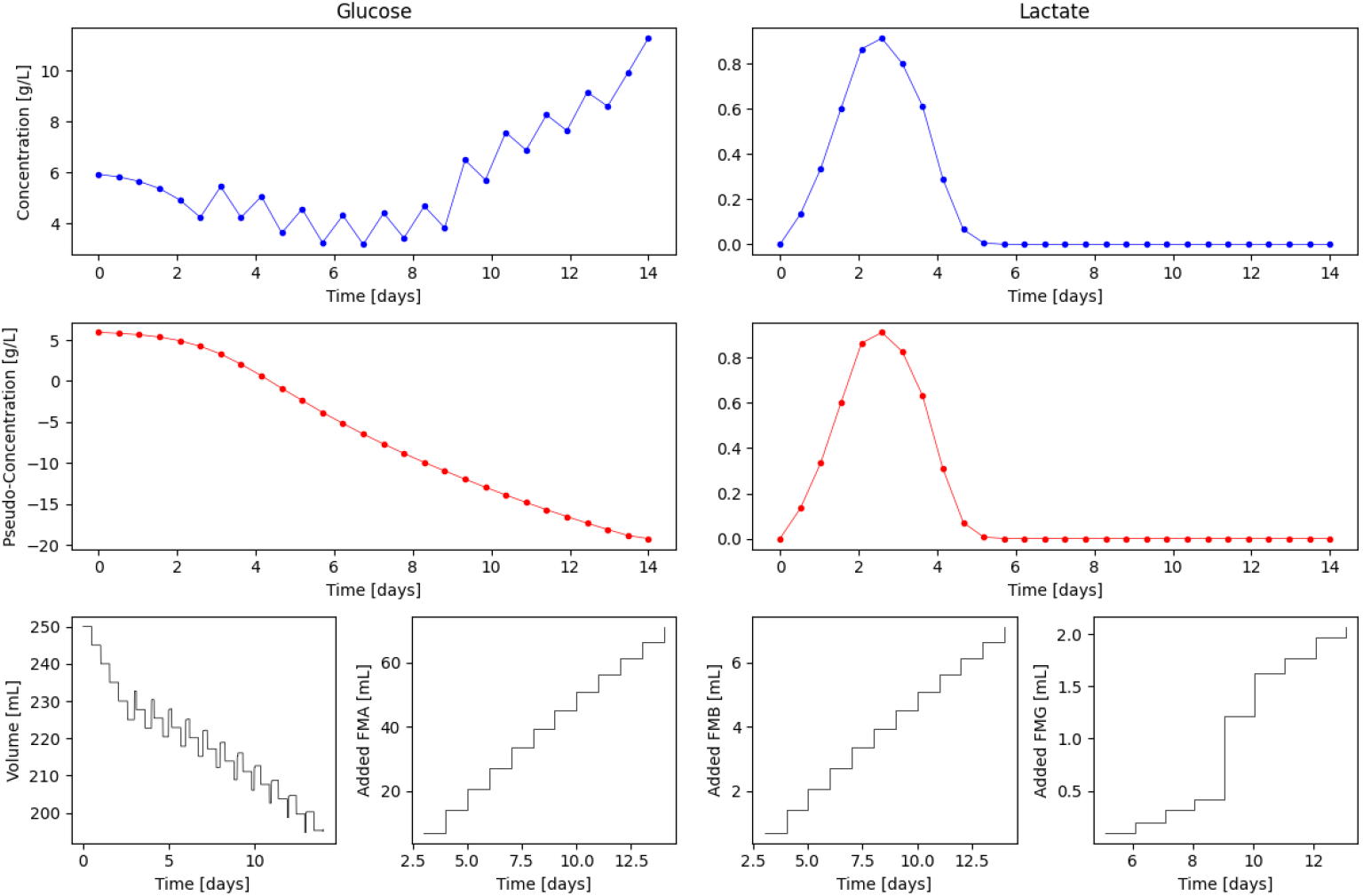
Example of a pseudo-concentration transformation from glucose and lactate concentration. Concentration data were generated using the model described in Section□2.7 under the following conditions: 28 uniformly spaced sampling points over the culture duration, each with a 5□mL sample volume, and no additional noise applied to the measured concentrations.

### Feature scaling

Scaling is applied to a set of observations 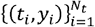, where *t* denotes time and *y* represents the observations by performing a general min-max normalisation. The operator 𝒯 is defined to transform the measurement values into the range [0, 1] using the following function:

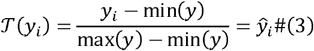

where 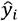 is the normalised value. To revert the normalised values to their original scale, the inverse transformation operator 𝒯 ^−1^ was defined as:

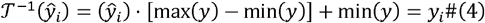

This scaling is applied out of convenience, allowing for arbitrary likelihoods, particularly when the likelihood function requires taking the logarithm. By normalising the data, compatibility with various likelihood functions is ensured, which enhances the flexibility and robustness of the analysis.

### Preprocessing summary

In summary, observations were first transformed using (1) and (2) to obtain pseudo-observations (denoted with the * symbol), which are then normalised by applying (3), resulting in normalised pseudo-concentrations (denoted with the ^ symbol):

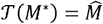

### 2.2. Prior Informed Interpolation Model

Assumptions on the behavior of metabolic rates (and hence the shape of metabolic trajectories) are applied by defining interpolation models as linear combinations of basis functions, referred to as the”mathematical model”. The goal is to interpolate a general set of points (observations) 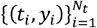,where *t* denotes time and *y* represents the observations (interpolated variable). To allow continuous interpolation between the points, we defined a linear combination of basis functions 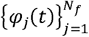 to approximate *y*(*t*):

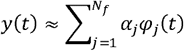

where *α*_*j*_ are the coefficients to be determined for each basis function *φ*_*j*_(*t*). For this work, the basis function takes the form of the integral of the sigmoid function:

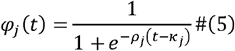

where *ρ* modulates function slopes, and denotes the center (allowing the basis function to be shifted by the direct distance *t* − *κ*). Note that other basis functions (i.e., gaussian and hyperbolic tangent) have been investigated (Andersson et al. 2025).

The integral of the sigmoid interpolation functions is fitted to the metabolite concentration data and then differentiated to obtain the rates. The initial point of interpolation is fixed by subtracting the difference between the initial point *y*_1_ and the sum of the basis functions evaluated at the initial time point *t*_1_. This leads to the following interpolation:

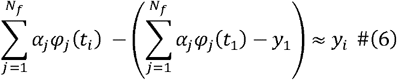

### 2.3. Specific Metabolic Rate

A continuous model for the metabolic rate *δ* of a metabolite was built by normalizing the rate of change of the metabolite concentration *M* with the viable cell density *X*:

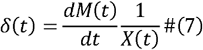

This was achieved by interpolating the normalized pseudo-density, 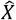, inserting the indefinite integral of the sigmoid basis (5):

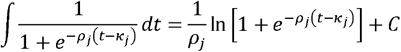

with a constant of integration *C* into equation (6) giving:

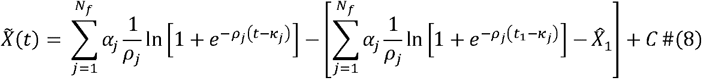

Furthermore, the distribution of basis center points was parameterized to a single variable by substituting *κ*_*j*_ with *g*_*j*_(*κ*) :

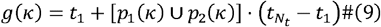

Here, *g*(*κ*) generates a vector of *N*_*f*_ center points distributed over the time interval 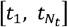, with one point for each basis function. Each of these points serves as the center of its respective basis function over the fitting interval (see Supplementary Materials for details).

This parameterization was introduced to reduce the degrees of freedom in the interpolation of metabolite trajectories. By using a single *κ* instead of individual *κ*_*j*_ values, the model becomes more parsimonious and computationally efficient. To maintain a generalizable framework, the number of basis functions was set according to the most complex metabolite profiles - an approach that, without dimensionality reduction, would result in an unmanageable number of parameters. The reduction scheme in Equation (9) addresses this challenge by preserving the necessary resolution for complex trajectories while keeping the model stable, interpretable, and compact.

By substituting the parameterized form given in (8) into (6), we obtain the interpolation for the normalized pseudo-concentration:

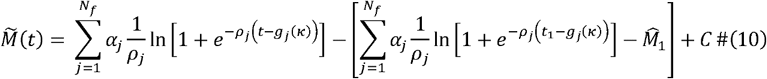

Finally, the parameterized derivative of (10), using the center point reduction scheme (9), is given by:

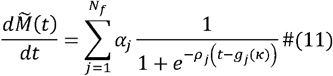

### 2.4. Statistical model

Two different models were used, one for the viable cell density and one for other metabolites.

#### Viable cell density

The available observations of viable cell density exhibited increasing variance with larger values, indicating a proportional relationship between the variance and observations. Hence, we used a log-normal distribution to model the observations. It was also assumed that there was an unknown constant measurement error for each observation, giving the following log-likelihood:

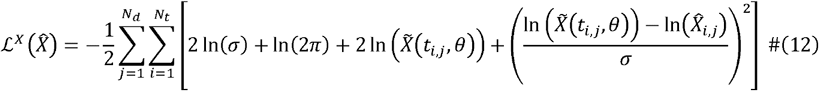

where 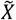 represents the model defined by equation (8), a noise parameter σ, and 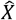 refers to a set of normalised measured pseudo-VCDs. The index *j* = 1,2,…,*N*_*d*_ allows the summation of several instances of observed trajectories for matching conditions. To enforce the positivity of the predicted viable cell density, the constraint 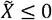 is incorporated in the mathematical model by dropping negative regressed values.

#### Metabolites

For metabolite observations, we assumed a normal distribution and unspecified measurement noise, giving the log-likelihood:

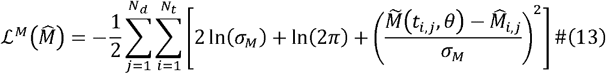

where 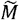 represents the model defined by equation (10), and 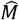 refers to a set of measured (and here also normalised) pseudo-concentrations. As before, the index *j* = 1,2,…,*N*_*d*_ allows the summation of several instances of observed trajectories. A single noise parameter σ was fitted for all measurements of a specific metabolite (i.e., *σ* = *σ*_*M*_). For a metabolite that is both produced and consumed, the base case in (13) applies (i.e., no additional constraints). For substrates (metabolites always consumed), we enforce a nonpositive rate of change 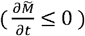; for products (metabolites always produced), we enforce a non-negative rate of change 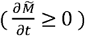. Again, the constraints are enforced by discarding results produced by the mathematical model outside the inequality constraints.

Model parameters are denoted by *θ* for n = 1,2,…,*N*_*θ*_ with *N*_*θ*_ being the total number of unique parameters per model. Uniform bounded priors were used for each parameter, 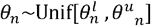, and a prior transform from uniformly distributed variables 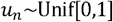 to the desired parameter ranges 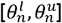 was applied using the following equation:

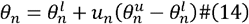

Finally, the measurement noise *σ* and *σ*_*M*_ were considered unknown parameters. We used a priori knowledge of the experimental environment to set the prior bounds for this parameter.

Having defined the models, we generate the posterior probability:

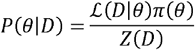

with Bayesian Inference using a model prior π, evidence *Z* and some data *D*. The evidence can be calculated by integrating the product of the likelihood and the prior over the entire parameter space *Ω*_*θ*_ as:

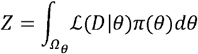

allowing the determination of the distribution parameters. For pseudo-density, the complete set of parameters are:

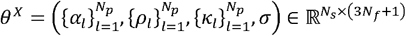

and for pseudo-concentrations:

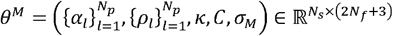

Once parameter distributions are inferred, the posterior predictive distribution for a new observation at an arbitrary timepoint 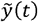 is defined as:

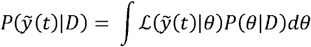

where 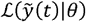 is the likelihood of the new observation given posterior parameters. We approximate the evidence using Nested Sampling, as implemented by Speagle (2020) and Koposov *et al*. (2024). Nested sampling estimates the evidence while simultaneously generating posterior parameter distributions. The algorithm iteratively replaces the least likely points in an initial set of samples with new points with higher likelihoods, maintaining a constraint on likelihood levels.

The complete procedure is summarised in Algorithm 1. First, instances of the mathematical models described in equations (8) and (10) are created. Subsequently, statistical models are constructed using the log-likelihood functions (12) and (13), and the prior transform (14). Data is transformed using equations (1) and (2), and then normalised according to equation (3). Then following parameter inference, samples from the posterior predictive distribution are transformed back to their original scale using the inverse of the min-max normalisation (4).

After normalisation pseudo-density can be approximated by 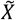 as:

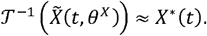

Given that 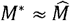 by minimizing (13) using chain rule expansion the rate of metabolic concentration can be defined as:

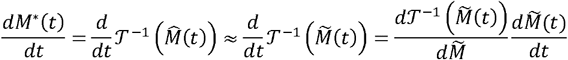

allowing the calculation of the cell-specific consumption rates, δ as in (7), repeated here in terms of pseudo observations for clarity:

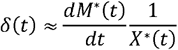

From the generated parameter distributions, trajectories can be sampled and generated for δ at arbitrary time points within the interpolated time interval.

In this work, we estimate VCD-normalized metabolic rates. Accordingly, for each experiment, a VCD model is fitted not only to obtain the growth rate but also to normalize the metabolic rates. Table 1 summarizes the prior settings for both metabolite and VCD models, highlighting the use of varying degrees of freedom. Metabolite and VCD measurements are assumed to have independent measurement and sampling errors. This assumption enables separate estimation and sampling of the parameter distributions. Without independence, the joint distribution of all parameters would need to be estimated, substantially increasing computational cost.

**Table 1.**
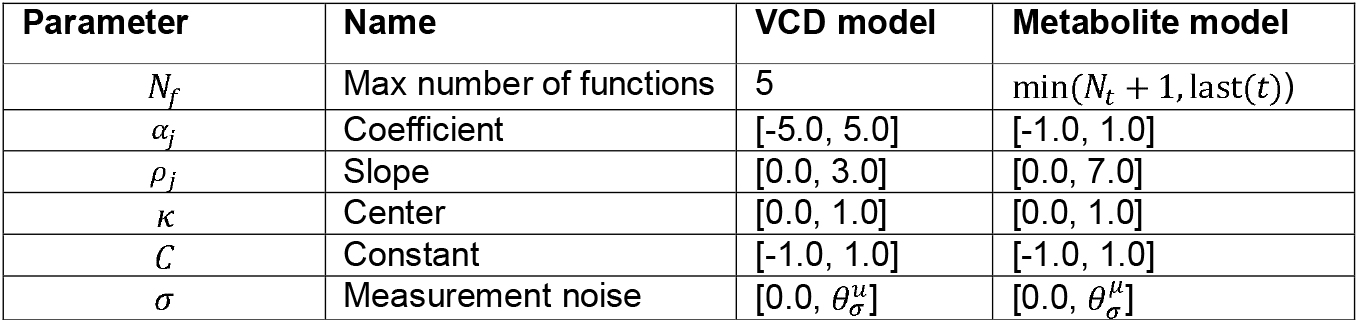
Parameter settings for the VCD and metabolite model.

##### Algorithm 1 Parameter Distribution and Rate Calculation

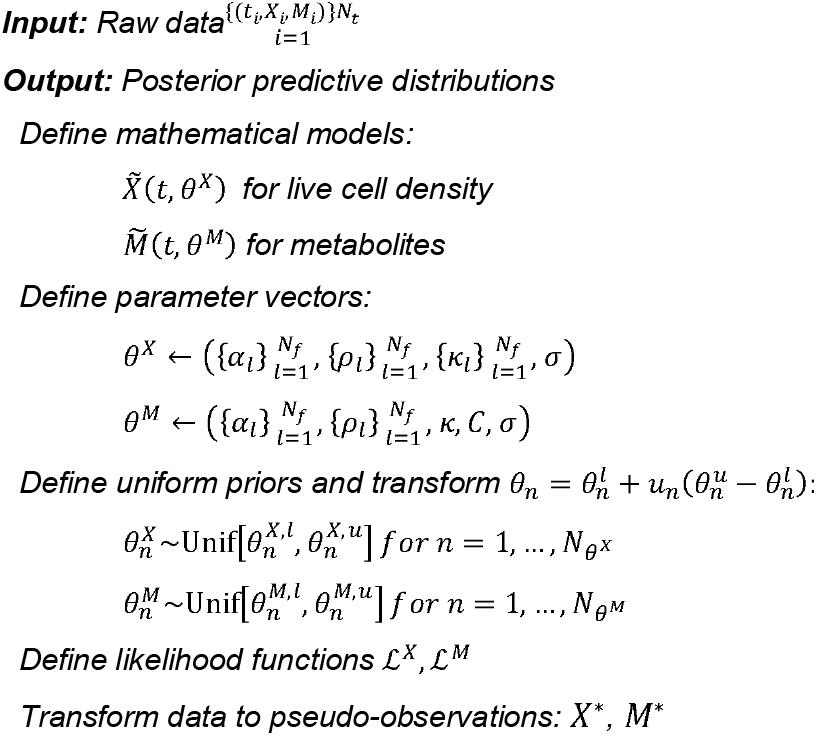
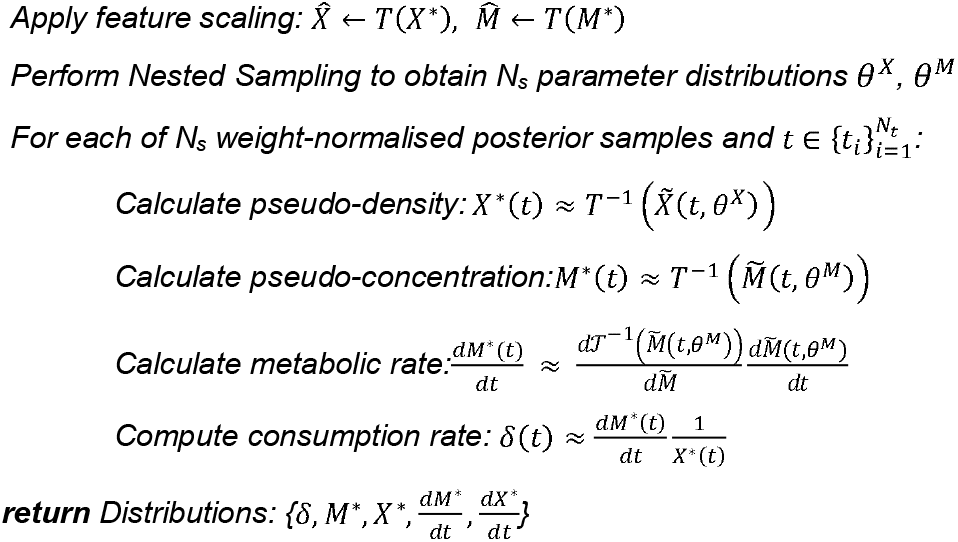

Parameter priors were specified as uniform distributions and tailored to the dataset under analysis. The limits chosen for the basis function slopes, for example, strongly affect the model’s ability to fit the data. Because the dataset is sparse and no fixed noise levels are imposed (the noise level is itself a fitted parameter), there is a competitive interplay between noise estimation and model flexibility in the likelihood. Higher assumed noise levels naturally expand model flexibility, as a broader range of models can then explain the data equally well. Finally, model center points (intercepts) were constrained to remain within the experimental time domain.

Prior bounds are typically chosen based on experience with similar experiments and the expected behavior of the system and data (e.g., constraining parameters to remain non-negative or adjusting the maximum slope of basis functions according to observed trends). These choices are then evaluated through visual inspection and Bayesian evidence. In many cases, users have reliable knowledge of the measurement noise (σ), in which case no freedom is required for this parameter. Practically, this is achieved by setting both the upper and lower bounds of σ to the known noise level. However, since precise noise levels are often unavailable, we adopt a more general formulation in which noise is treated as a free parameter. To support this, we provide heuristic strategies for defining suitable prior bounds on the noise level.

### 2.5. Model selection framework

Nested sampling is well-suited for Bayesian inference because it not only estimates model parameters but also calculates Bayesian evidence, enabling rigorous model comparison based on their ability to explain the observed data. By defining a family of models with varying characteristics, we can evaluate them through the log-evidence metric (log(Z)). Algorithm 2 explores model configurations by iterating over a specified range of degrees of freedom, pairing each with different interpolation models. Every combination is instantiated and passed to the sampler, which applies nested sampling to estimate parameters and evidence. Models that fail to converge (e.g., yielding NaN, infinite negative, or empty values) are discarded, and the remaining models are ranked by their log-evidence. The top-ranked subset can then be selected for detailed analysis

#### Algorithm 2 Model configuration, sampling, and ranking

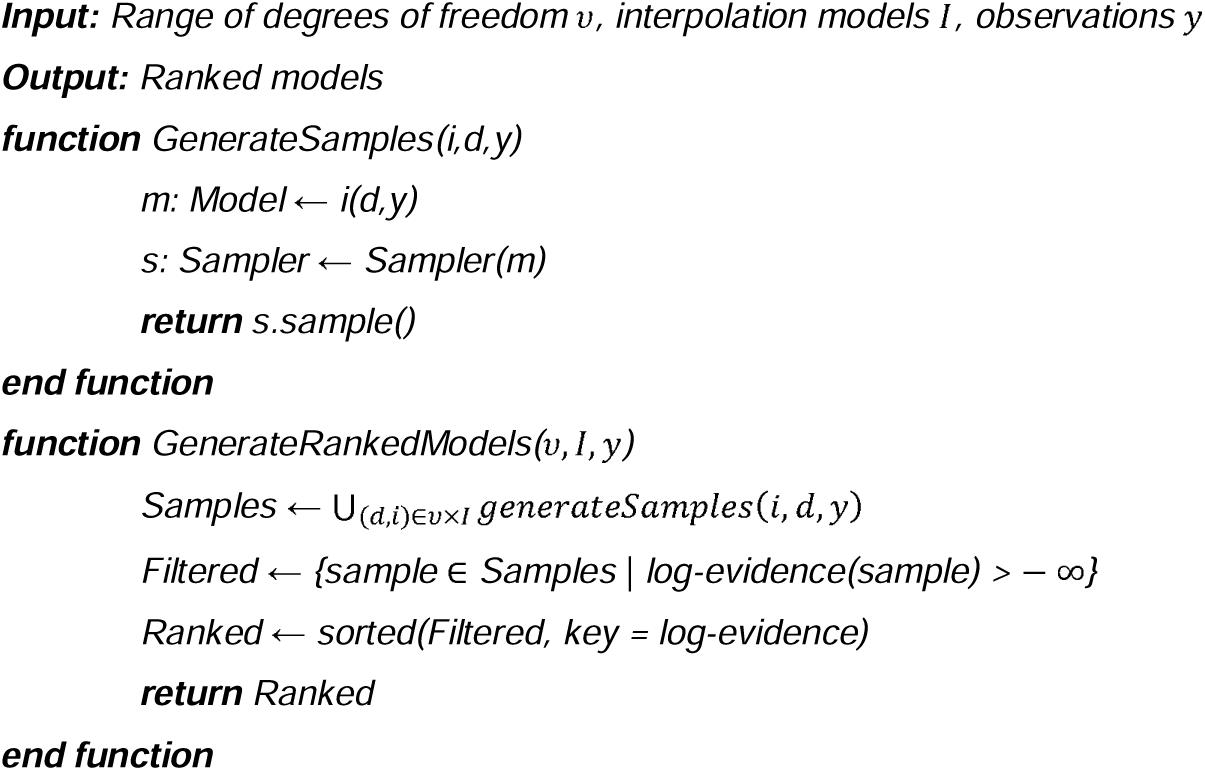

In Algorithm 2, the parameter *v* (degrees of freedom) corresponds to the number of basis functions used to model the data. This value can be specified based on prior knowledge - for example, if the data suggest the presence of *n* distinct growth phases, then *v* = *n* may be a suitable choice. Alternatively, a model selection procedure can be employed: the number of basis functions is varied from 1 up to a reasonable maximum (e.g., the total number of days *d*, under the assumption that more than one growth phase per day is unlikely), and the optimal value is chosen based on the highest Bayesian evidence.

### 2.6. In silico data generation

To evaluate the rate estimation framework proposed in this study, we developed a macroscopic mechanistic model of CHO cell metabolism and bioprocess dynamics. The model is formulated as a system of ordinary differential equations (ODEs) that describe viable cell growth, cell death, product formation, and the metabolism of key nutrients and byproducts - namely glucose and lactate. This type of structured macro-kinetic modeling has been widely used in the bioprocess field to simulate fed-batch dynamics (Sha et al., 2018; Richelle et al., 2022; Bayer et al., 2023). The model equations are as follows:

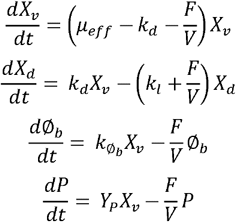

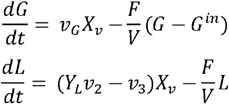

where *X*_*v*_ is the viable cell density (VCD, 10^6^ cells/mL), *X*_*d*_ is the concentration of dead cells (10^6^cells/mL), ∅_*b*_ is the “biomaterial” concentration (g/L) representing accumulated byproducts (e.g., ammonia, host cell proteins) as a lumped variable (Richelle et al., 2022), *P* is the concentration of the protein product synthesized by the cells (mg/L), *G* and *G*^*in*^ are, respectively, the glucose concentration in the culture medium and the feeding (g/L) and *L* is the lactate concentration (g/L). *F* is the feed flow rate of the feed (L/day), and *V* is the bioreactor volume (L). *µ*_*eff*_ is the effective growth rate (day^-1^), *k*_*d*_ is the death rate constant (day^-1^), *k*_*l*_ is the lysis rate constant (day^-1^), 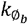 is the biomaterial secretion rate constant (set to 1 to ensure identifiability). *Y*_*P*_ and *Y*_*L*_ are pseudo-stoichiometric coefficients for product and lactate formation, respectively.

The glucose uptake rate *v*_*G*_ is defined using a saturation-type kinetics modulated by inhibitory effects from biomaterial accumulation:

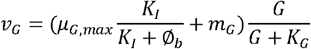

Overflow and aerobic metabolism are split using the following definitions:

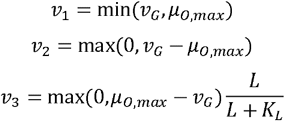

The cell growth rate *µ*_*eff*_ is represented as a weighted sum of the metabolic modes:

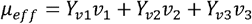

This hybrid formulation allows the model to represent transitions between aerobic and anaerobic metabolism (e.g., glucose overflow leading to lactate accumulation), while enabling the simulation of feeding strategies and process dynamics. In silico datasets generated using this model provide a controlled, reproducible setting to benchmark metabolic rate inference methods under varying noise and sampling conditions. The parameter values and process information used to generate the in silico dataset (see Supplementary Materials) were selected to reflect the variability typically observed in real experimental datasets (data not shown).

## 3. Results

### 3.1. In-silico bioprocess simulation and dataset generation

A dynamic CHO fed-batch model (Section 2.7) was used to generate realistic process trajectories under varying sampling and noise conditions, enabling a controlled benchmark for evaluating rate estimation methods. To assess the robustness of the proposed rate estimation method, multiple in-silico datasets were generated by varying three experimental design factors (total of 48 datasets generated):

1. **Number of samples:** Three configurations were tested with 7, 14, or 28 sampling points, uniformly distributed over a 14-day culture period.
2. **Sample volume:** At each sampling time, four different sample volumes were considered (2.5, 5.0, 7.5, and 10.0□mL).
3. **Noise level:** Gaussian noise was applied to each measurement, proportional to its magnitude, with four different levels: level□1 (□=□0; 0%□CV), level□2 (□=□0.05; 5%□CV), level□3 (□=□0.10; 10%□CV), and level□4 (□=□0.15; 15%□CV). Noise was applied independently to each measured variable following:

where is the simulated measurement from the model and is the selected noise level.

The overall workflow for the *in silico* bioprocess simulation is summarized in Figure□3a. The procedure begins with the specification of a dynamic CHO fed-batch model and the definition of user-selected sampling volumes, which determine the rate of culture volume decrease over time. The model, in combination with the imposed volume trajectory, is then used to simulate continuous “true” trajectories for all process variables. From these trajectories, the user-specified number of sampling points is extracted, after which Gaussian noise is applied according to the chosen noise level. The final output is a fully synthetic concentration dataset that integrates the combined effects of the selected experimental design parameters.

**Fig□3.**
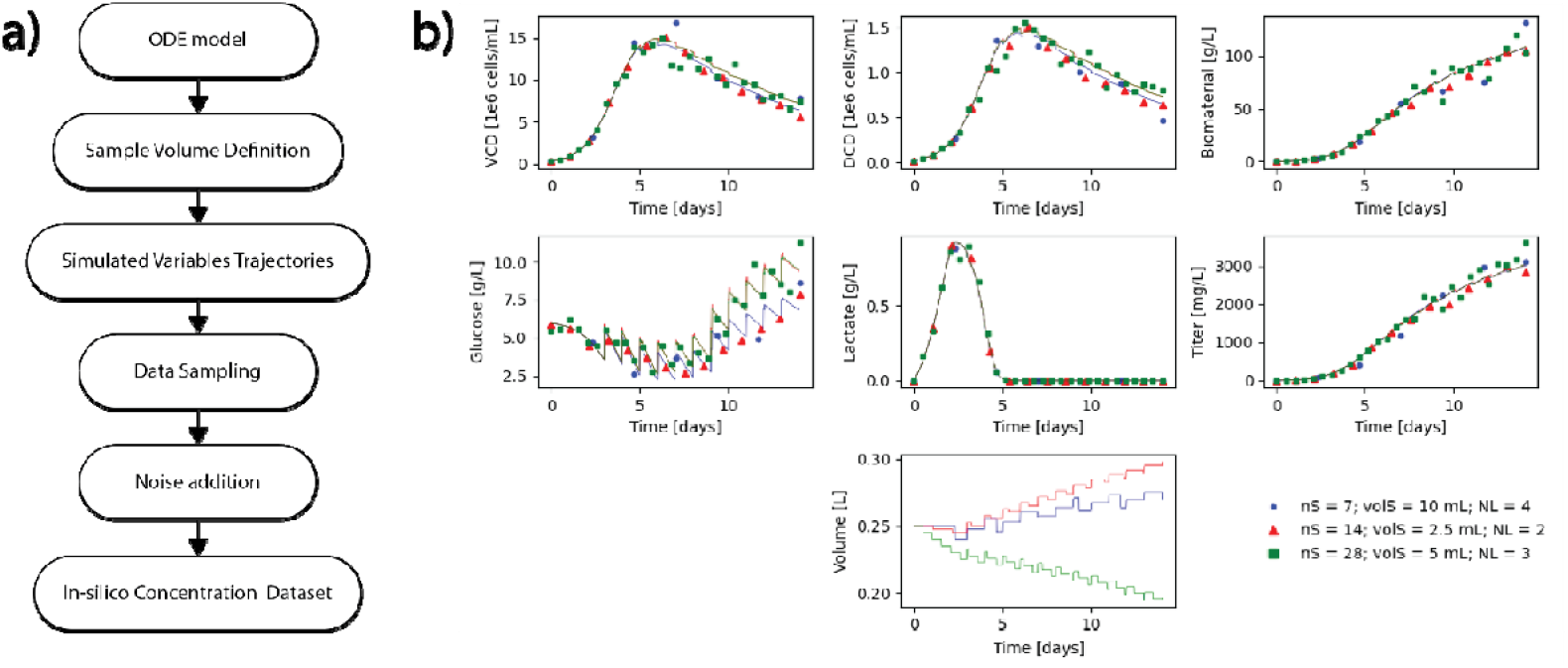
Workflow for *in silico* bioprocess simulation and example results. a) Stepwise procedure from ODE-based process modeling to generation of noisy, down sampled datasets. B) Simulated variable trajectories for three design configurations (nS = number of samples, volS = volume of the samples, and NL = noise level). Sampling volume affects glucose concentration most strongly, with minor effects on VCD and DCD, while lactate, titer, and biomaterial remain largely unaffected.

Figure□3b presents example trajectories for the model variables under three distinct experimental design scenarios. While the simulated lactate, titer, and biomaterial profiles remain largely unaffected by changes in sampling volume, glucose concentration displays pronounced variations resulting from the interaction between feeding events and volume removal. The viable cell density (VCD) and dead cell density (DCD) trajectories also exhibit moderate differences associated with sampling volume, particularly during the late exponential and decline phases. These results demonstrate that certain process variables are more sensitive to sampling volume definitions than others.

### 3.2. Bayesian–nested sampling method for rate estimation and concentration reconstruction

The Bayesian–nested sampling framework enables accurate and well-characterized estimation of metabolic rates from sparse and noise-free or noisy measurements. Figure□4 presents representative results obtained using the MetRaC procedure applied to synthetic in silico data generated under ideal conditions - daily sampling of 5□mL with no added noise.

**Fig 4.**
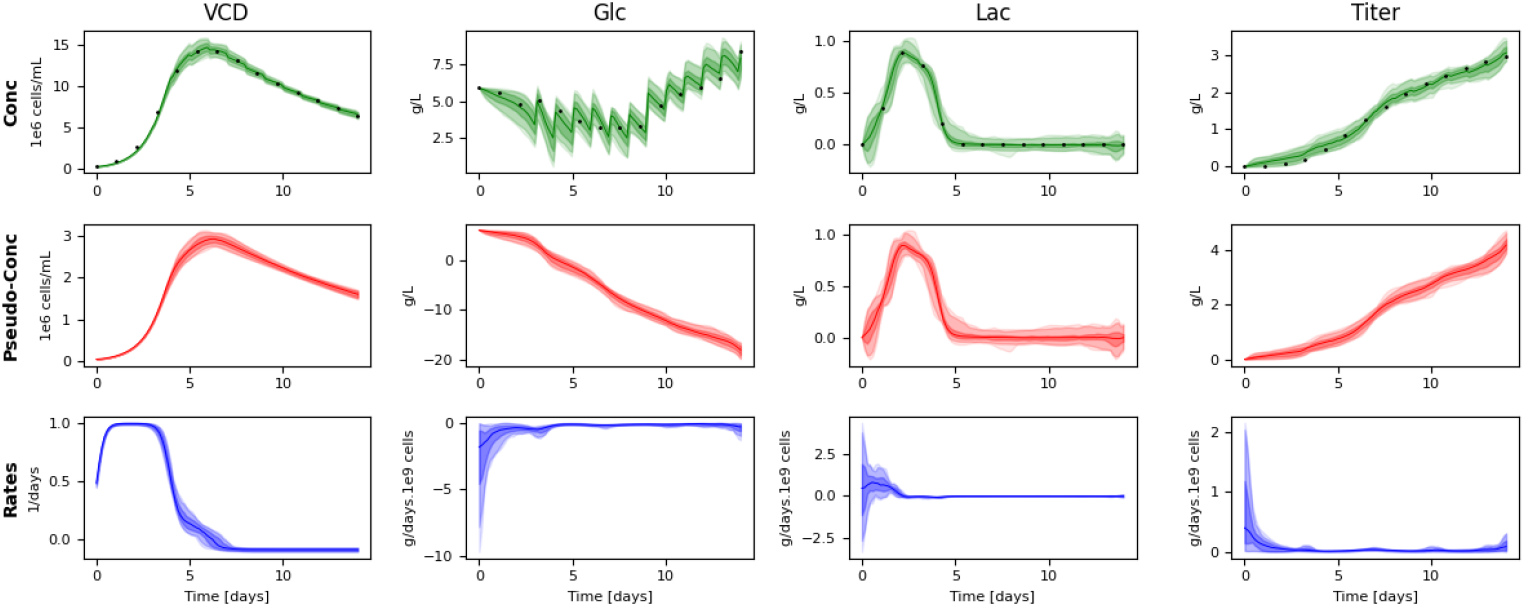
Posterior predictive distributions generated by the MetRaC framework from in silico data with daily sampling (5□mL) and no added noise. Each column corresponds to a process variable: viable cell density (VCD), glucose (Glc), lactate (Lac), and titer. The top row shows the reconstructed concentration profiles with black dots representing the synthetic measurement data. The second row shows the inferred pseudo-concentration distributions, and the third row displays the corresponding metabolic rate posteriors. Shaded areas represent 68%, 95%, and 99% confidence intervals; colored lines denote the posterior medians.

The framework infers a posterior distribution over the pseudo-concentration profile (Figure□4, second row), which is used to fit an interpolation model. This posterior is then propagated through the metabolic model to yield a posterior distribution over estimated rates (Figure□4, third row). Additionally, we back-propagate the pseudo-concentration posterior to reconstruct the original concentrations, capturing the variability “perceived” by the method due to the finite sampling volume and temporal sparsity (Figure□4, top row).

Even in the absence of measurement noise, the results reveal that data sparsity alone can induce noticeable uncertainty in the inferred concentration and rate profiles. This underscores the value of the Bayesian–Nested sampling approach in providing robust estimates of metabolic rates, even under ideal experimental conditions.

The metabolic rate estimates obtained using MetRaC can be used to reconstruct concentration trajectories by integrating them into a generic ordinary differential equation (ODE) system describing the fed-batch bioprocess dynamics:

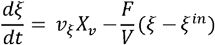

where denotes the state variable of interest (e.g., VCD, glucose, lactate, or titer), represents the specific rate estimated by MetRaC, is the viable cell density (VCD), is the feed rate, is the culture volume, and is the concentration of in the feed.

This forward simulation approach (Figure 5a) differs from the back-propagation scheme shown in Figure□4, as it integrates the estimated rates over time, introducing a different propagation of uncertainty stemming from noise, data sparsity, and volume dilution effects (see also Supplementary Figure□S1).

**Fig 5.**
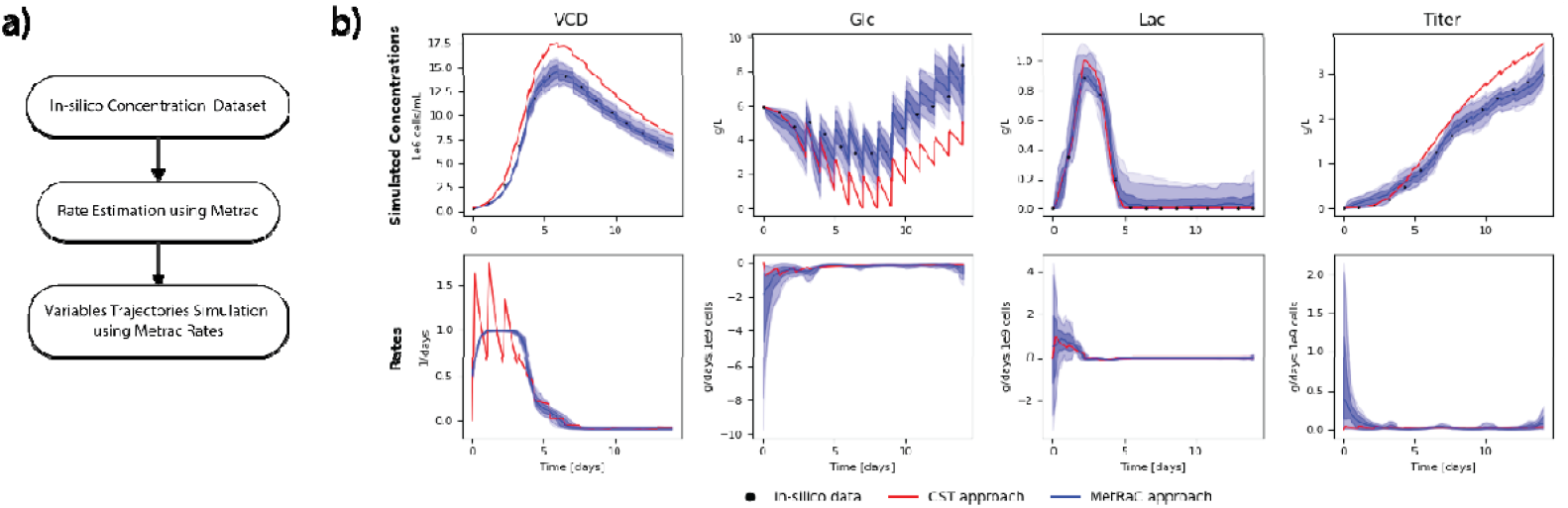
(a) Overview of the workflow for concentration trajectory simulation using MetRaC rate estimates. Simulated in-silico concentration data are used to estimate metabolic rates via MetRaC, which are then integrated into a bioprocess ODE model to reconstruct state variables over time. (b) Comparison of simulated concentration and rate trajectories for VCD, glucose, lactate, and titer using MetRaC (blue) versus a standard CST rate approximation (red). Shaded areas represent 68%, 95% and 99% credible intervals, and black dots indicate in-silico data.

Figure□5b compares the concentration simulations obtained using MetRaC-based rate estimates with those derived from a commonly used constant-rate approximation (further referred to in the text as “CST approach”). This reference method assumes a linear change in metabolite concentration between consecutive measurements, producing a piecewise constant estimate of the specific rate. Specifically, the average rate of change is computed as the ratio of concentration differences to time intervals between measurements. To obtain specific consumption rates, these average rates are normalized by viable cell density (VCD), which is interpolated using a log-linear scheme. For time points, the VCD is approximated as:

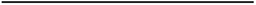

where and are the VCD measurements at times and, respectively. The exponential of this interpolated log-VCD is then used to normalize the constant rate estimate.

While this CST rate method provides a simple and widely adopted baseline, it does not account for uncertainty in measurements nor the sparsity of the data. In contrast, the MetRaC approach incorporates uncertainty quantification and offers smoother, more consistent reconstructions. As shown in Figure□5b, the discrepancies in estimated rates between the two approaches propagate into substantial differences in simulated concentration profiles - highlighting the importance of uncertainty-aware methods for robust process modeling.

### 3.3. Impact of sampling strategy and noise level on rate estimation accuracy and reconstruction quality

To quantify the impact of experimental design parameters on rate estimation accuracy and concentration reconstruction quality, we examined the effects of sample number (*nS*), sample volume (*volS*), and measurement noise level (*NL*) on a range of RMSE metrics. Normalized RMSE values were computed by comparing predictions against the “true” in-silico simulation outputs, scaled by the maximum signal amplitude for each variable. The relationship between RMSE and the three experimental factors was assessed using Random Forest regression, with feature importance expressed as the mean decrease in impurity (Gini importance) and reported as the percentage of variance explained (Figure 6).

**Fig 6.**
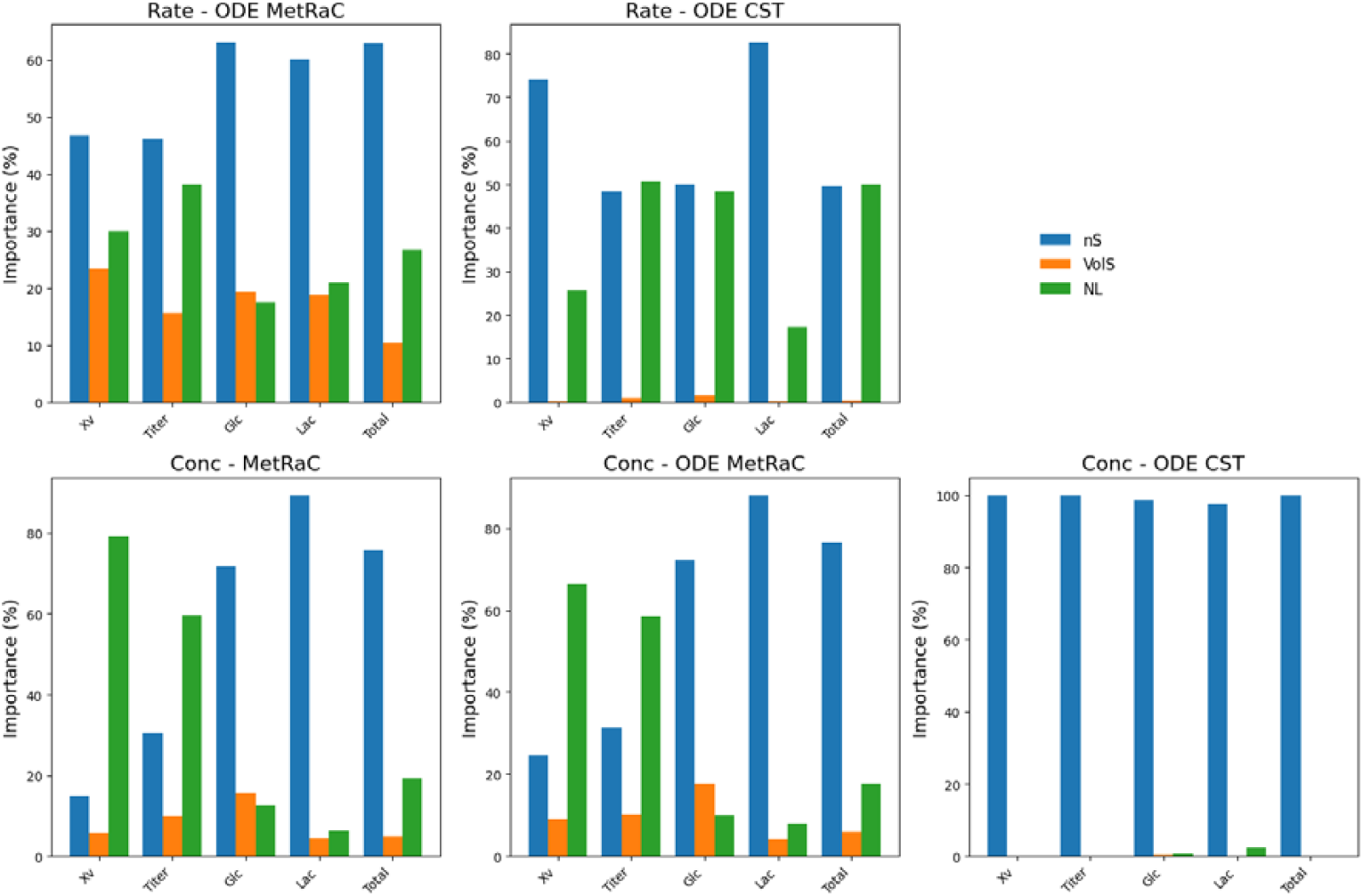
Relative importance of experimental design parameters on RMSE for rate estimation and concentration reconstruction. Importance was quantified using a Random Forest regression model (Gini importance) and expressed as the percentage of variance explained for each factor: sampling frequency (*nS*), sampling volume (*VolS*), and noise level (*NL*). Panels show results for: rate estimation using MetRaC (top left) and constant rate (top middle), concentration reconstruction via MetRaC backpropagation (bottom left), ODE simulation with MetRaC rates (bottom middle), and ODE simulation with constant rates (bottom right).

Across all configurations, *nS* consistently emerged as the most influential factor for rate estimation, typically accounting for 40–60% of the explained variance. This effect was especially pronounced for ODE-based methods (both MetRaC and CST), where sparse sampling substantially increased rate RMSE. *NL* was the second most influential parameter overall, but in certain concentration reconstruction cases - particularly with CST approach - it exceeded the importance of *nS*, indicating that measurement precision can, in some cases, outweigh temporal resolution in determining concentration accuracy. *VolS* generally had a minor role, with importance values below 15%, though it became more relevant for lactate and glucose concentrations in MetRaC, where lower sample volumes were consistently associated with higher RMSE.

Importantly, the ranking of factors differed between estimation and reconstruction tasks. In MetRaC backpropagation reconstructions, *NL* often dominated - except for lactate, where *nS* prevailed - whereas ODE-based reconstructions depended almost entirely on *nS*, with negligible influence from the other factors. These findings were robust to the choice of importance metric: repeating the analysis with a permutation-based approach (Figure S2) yielded consistent conclusions.

These patterns were mirrored in the heatmaps of normalized RMSE across parameter combinations (Figure 7). Lower *NL* and higher *nS* led to smaller RMSEs for both rates and concentrations, although the magnitude of improvement varied between the MetRaC and CST configurations. Increasing *VolS* reduced errors in some cases, but the effect was generally weaker and less systematic compared to *nS* and *NL*.

**Fig 7.**
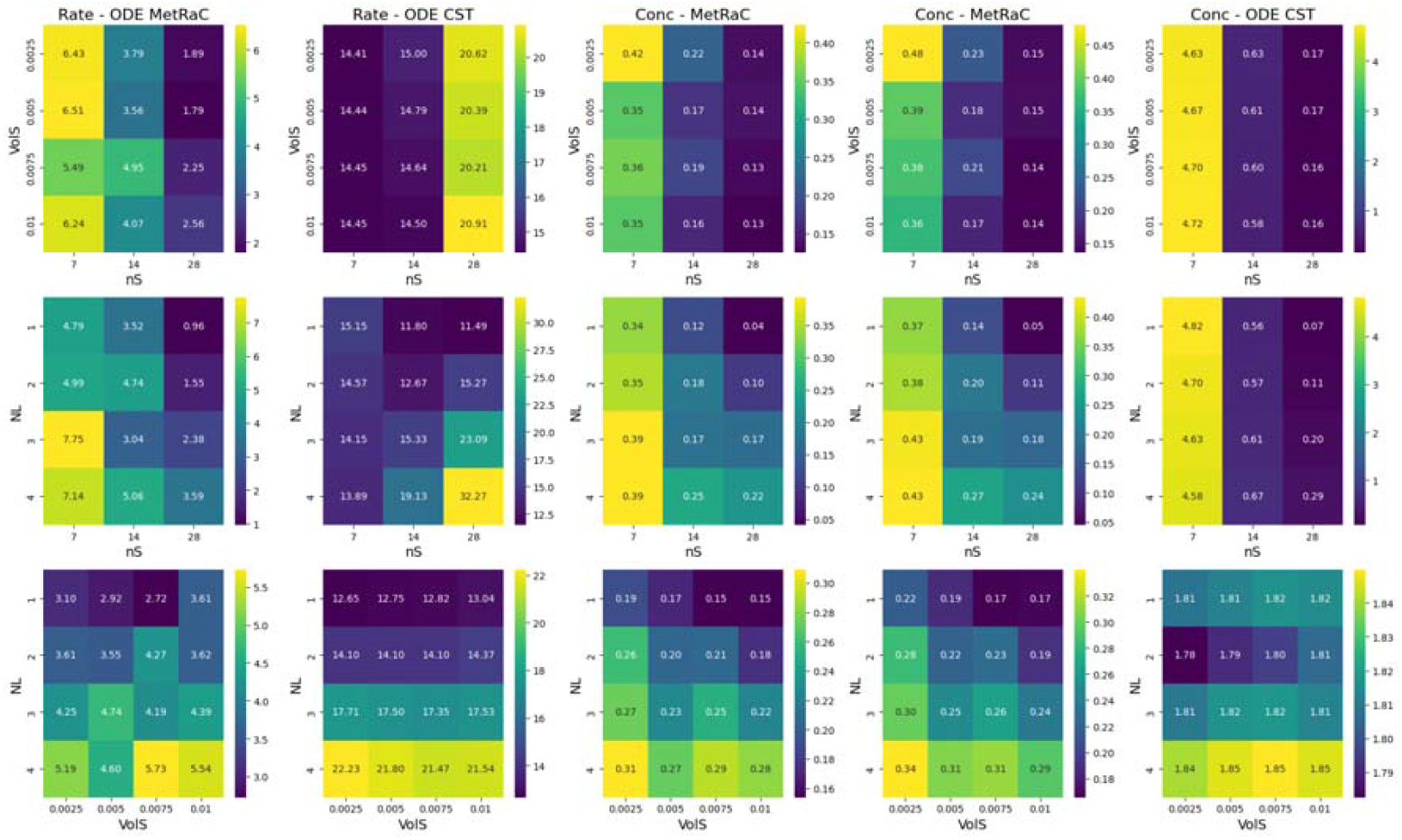
Heatmaps of normalized RMSE values for rate and concentration predictions across combinations of *nS, VolS*, and *NL*. Results are displayed for: rate estimation using MetRaC (1^st^ column), constant rate (2^nd^ column), concentration reconstruction via MetRaC backpropagation (3^rd^ column), ODE simulation with MetRaC rates (4^th^ column), and ODE simulation with constant rates (5^th^ column).

Finally, we examined the relationship between total rate RMSE and total concentration RMSE across all parameter combinations (Figure 8). For the MetRaC configuration, a clear positive correlation emerged (Pearson *r* ≈ 0.8): conditions yielding lower rate errors also produced more accurate concentration reconstructions. In contrast, CST results displayed a more clustered pattern, with certain combinations achieving low concentration errors despite moderate to high rate RMSEs.

**Fig 8.**
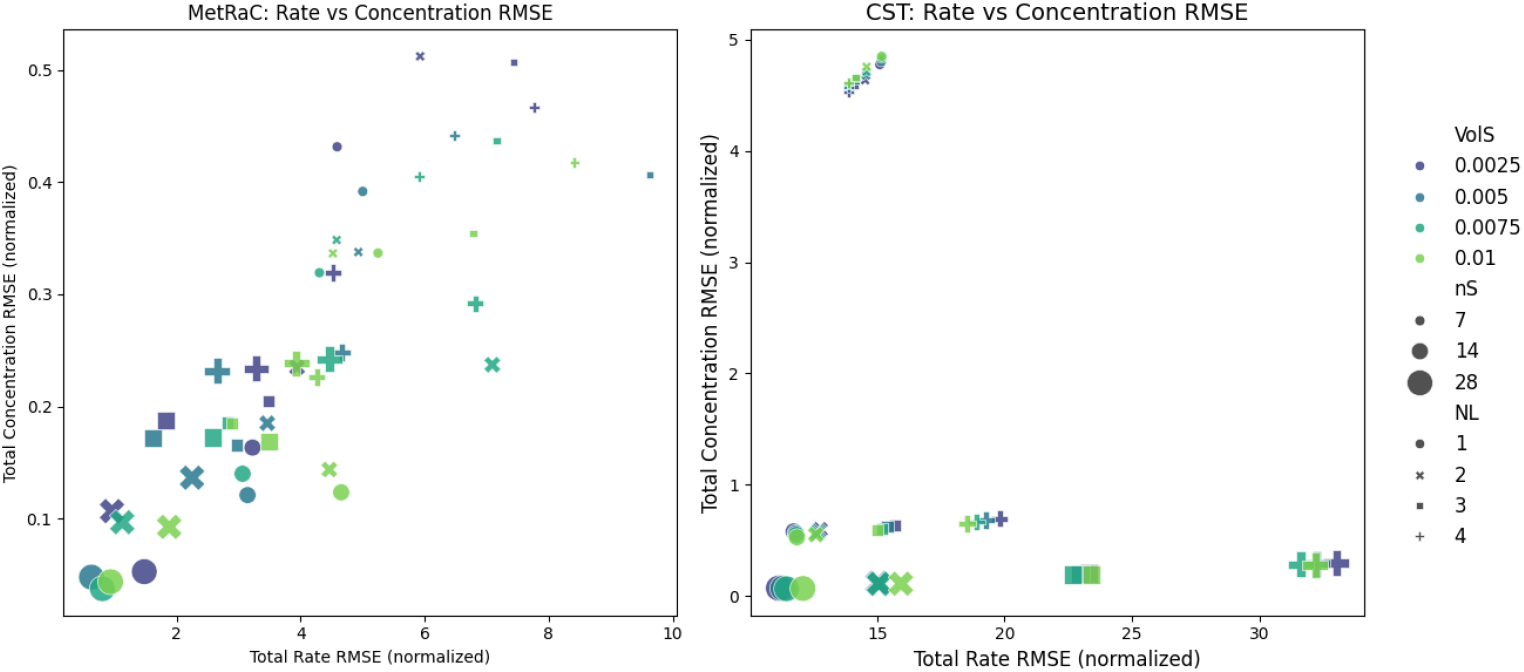
Scatterplots comparing total rate RMSE (x-axis) and total concentration RMSE (y-axis) for the MetRaC and CST configurations. Marker color encodes *VolS*, marker size encodes *nS*, and marker shape encodes *NL*.

Overall, these results highlight that while temporal resolution, sample volume, and measurement noise all influence rate and concentration accuracy, their impact is not uniform across estimation approaches. The Bayesian MetRaC method consistently demonstrated lower sensitivity to adverse experimental conditions than the constant-rate (CST) approach, maintaining comparatively high accuracy even under sparse sampling, low sample volumes, or elevated noise. Although both methods benefited from improved temporal resolution and reduced measurement noise, CST’s performance degraded more sharply when these parameters were suboptimal, reflecting its stronger dependence on high-quality, high-frequency data. In contrast, MetRaC’s probabilistic framework provided greater robustness, enabling it to retain predictive power and more faithfully reproduce process trajectories across a wider range of experimental scenarios.

## 4. Discussion

In this study, we have developed a probabilistic framework for bioprocess modeling that addresses the limitations of traditional rate calculations from sparse concentration data. Our approach leverages Bayesian inference and Nested Sampling to provide robust estimates of metabolic rates, enabling better understanding and modeling of bioprocesses. The MetRaC framework transforms sparse and noisy concentration measurements into continuous rate profiles, incorporating biological insights directly into the model structure. This probabilistic approach offers several advantages, including robustness to noise, interpretability, and the ability to quantify uncertainty.

The proposed framework enhances bioprocess understanding by providing accurate and reliable estimates of metabolic rates, even under challenging experimental conditions characterized by data sparsity and measurement noise. The ability to quantify uncertainty in rate estimates is particularly valuable for risk assessment and process validation, ensuring that predictions are both informative and reliable.

Despite its advantages, the framework has certain limitations. The computational cost associated with Bayesian inference and Nested Sampling can be substantial, particularly for high-dimensional parameter spaces. Additionally, the sensitivity of the model to prior assumptions requires careful consideration and domain expertise to ensure accurate parameter estimation. The framework also demands comprehensive data requirements, including well-characterized prior distributions and sufficient sampling frequency to capture dynamic transitions in biological systems.

While the current implementation focuses on retrospective rate estimation, the Bayesian formulation of our framework naturally lends itself to future extensions for real-time control and near-horizon predictive modeling. In such an application, newly acquired process data could be assimilated in an at-line fashion to continuously update posterior rate estimates, while preserving quantified uncertainty. These updated estimates could then feed into process simulations or model predictive control schemes to forecast system behavior over short horizons, enabling proactive and informed process adjustments. The probabilistic nature of the Bayesian approach ensures that both predictions and control actions would be accompanied by credible intervals, allowing operators to weigh performance gains against associated risks - an advantage particularly relevant for decision-making under uncertainty in industrial bioprocesses.

Future work will focus on translating this potential into practice by integrating the probabilistic framework with control systems to enable real-time process monitoring and optimization. Hybrid modeling approaches that combine mechanistic and data-driven components may also be explored to improve scalability and computational efficiency. Additionally, targeted algorithmic improvements and parallelization strategies can be pursued to reduce computational costs, making the approach more suitable for deployment in time-sensitive industrial settings.

Overall, the probabilistic framework presented in this study represents a significant advancement in bioprocess modeling, offering a robust and interpretable tool for estimating metabolic rates from sparse and noisy data. By addressing the limitations of traditional rate calculations, our approach paves the way for more accurate and reliable bioprocess models, with potential applications in various fields, including biotechnology, bioengineering, and chemical engineering.

## Acknowledgements

We thank Shanti Pijeaud and others in the Sartorius Corporate Research team for their support in the development of this study.

