## Supplementary Materials for "Overcoming the Limits of Traditional Rate Calculations from Sparse Concentration Data: A Probabilistic Framework for Bioprocess Modeling"

#### *Pseudo-concentration calculation for perfusion processes*

The pseudo-transformation calculations described in the main manuscript require adaptation for perfusion processes. The evolution of the metabolite concentration $[met_i]$, (for $i = 1, \ldots, N_m$ metabolites – index $i$ is omitted for readability) is governed by the following equations:

$$\frac{dX}{dt} = X(t)\left(\mu + \frac{F_h(t) - F_{in}(t)}{V(t)}\right) \quad (S1)$$

$$\frac{d[met](t)}{dt} = \delta(t)X(t) + \frac{F_h(t)}{V(t)}\left([met]_{in} - [met](t)\right) \quad (S2)$$

$$V(t = T) = V_0 + \int_0^T \left(F_{in}(t) - F_b(t) - F_h(t)\right)dt \quad (S3)$$

where $X$ is the viable cell density, $F_h$ is the harvest flow, $F_{in}$ is the feed flow, and $F_b$ is the bleed flow, and$[met]_{in}$is the concentration of the metabolite in the feed flow. To find the function $\delta(t)$ describing the specific metabolic rate in (S2), equation (2) is integrated:

$$\int \frac{d[met](t)}{dt}dt = [met](t) = \int \delta(t)X(t)dt + \int \frac{F_{in}(t)}{V(t)}\left([met]_{in} - [met](t)\right)dt \quad (S4)$$

To account for the second term on the right-hand side (the feeding effect), a spline is fitted to [met]($t$), and this is divided by the VCD to get $\delta(t)$.

In cases where only bolus feeds are present, without any bleeding or harvesting (as in fed-batch culture operations), the approach using the ADF as described in the main manuscript can be applied to eliminate the second term. This is because only the concentration and feed volume at the time of feeding are required. However, if continuous feeding, bleeding, or harvesting is involved (continuous culture operations), removing the second term from the equation becomes significantly more challenging. To understand why, note that:

$$\int \frac{F_{in}(t)}{V(t)}\left([met]_{in} - [met](t)\right)dt \quad (S5)$$

can be removed since $[met](t)$ only needs to be evaluated at the time of feeding and at the point of measurement values. It becomes increasingly difficult to know the concentration without the feed if only the concentration fed is known, but not the concentration with feeding. This is the case for a continuous flux: at each time point, the change in metabolite concentration depends on the specific metabolic rate and the current concentration. To solve this, either the differential equation needs to be solved as part of fitting the spline, or assumptions need to be made on the behavior of $[met](t)$.

Solving the differential equation could be done by fitting a spline to $\delta(t)$, calculating the value of $[met](t)$, and using the difference between the calculated and measured concentrations to find the best representation of $\delta(t)$. However, this is likely computationally expensive. The issue is that we are trying to find $\delta(t)$, which is part of the derivative (i.e., it is a term of equation (S2)), while we only have access to the measured data ($[met](t)$), which is the integrated solution of the derivatives (including $\delta(t)$) as set out in equation (S4). One possible approach to solve this is to guess a functional form for $\delta(t)$ (and all other unknown parts of the system), then estimate all necessary parameters in the system by solving the ODE for those functions and parameters, and comparing the result to the measured data. Essentially, this fits everything in the system at once.

In the second approach, we assume that the metabolite concentration is linear between measurement points, which is a reasonable assumption if the measurements are frequent enough. $[met](t)$ can then be replaced with the linear approximation $L^{met}(t)$ on the interval. As feeding is usually done by setting a constant rate of flux, it can be assumed that $F_{in}$ is a constant, and $V(t)$ is a piecewise linear function of time. Then:

$$\int \frac{F_v(t)}{L^V(t)}\left([met]_{in} - L^{met}(t)\right) dt \qquad (S6)$$

can simply be evaluated. From this point, the analysis can proceed as in the bolus case. Specifically, if we can remove the second integral on the right-hand side in equation (S4), then the remaining problem is equivalent to the bolus case, where we could account for the feeding by doing the ADF and removal of mass. Removing the feeds by solving a simpler integral would give the equivalent to the pseudo-concentration of the bolus case.

#### ***Basis center point reduction.***

The sequence $g(\kappa)$ (15) is split left and right of $\kappa$ at the index $N_p = \left\lceil\frac{N_f+1}{2}\right\rceil$ into the sets $p_1$ and $p_2$ which make up a piecewise sinusoidal distribution (i.e., on each side of the time point parametrised by the single parameter $\kappa$, we get two similar distributions of points generated by a sinusoidal function, where the distributions of points either side of the time point defined by $\kappa$ are squeezed into two different periods, one comprising all $p_1$ points before the value of $\kappa$, and one comprising all $p_2$ points after $\kappa$), such that a single parameter $\kappa$ needs to be fitted for all basis functions in a model. The set $p_1$ is bounded by [0, $\kappa$]:

$$p_1(\kappa) = \left\{\frac{\kappa}{2}\left[\sin\left(s_1 - \frac{\pi}{2}\right) + 1\right]\right\} \qquad (S7)$$

$$s_1 = \left\{\frac{n\pi}{N_p - 1}\middle| n = 0,1,2,\ldots,N_p - 1\right\} \qquad (S8)$$

and $p_2$ by ($\kappa$,1]:

$$p_2(\kappa): \left\{\frac{\kappa + (1-\kappa)}{2}\left[\sin\left(s_2 - \frac{\pi}{2}\right) + 1\right]\right\} \qquad (S9)$$

$$s_2 = \left\{\frac{n\pi}{N_f - N_p + 1}\middle| n = 1,2,\ldots,N_f - N_p\right\} \qquad (S10)$$

The procedure essentially clusters the basis center points with a diminishing spacing around one chosen point along $t$. For example, with a $\kappa$ corresponding to a time point in the middle of the time course, this creates basis center points at intervals of diminishing size from $t = 0$ to $\kappa$ then increasing in size from $\kappa$ to the end of the time course. By contrast, with a $\kappa$ corresponding to a time point at the start of the time course, this creates basis center points at intervals of increasing size to the end of the time course.

Alternative strategies are possible to deal with the parameters $\kappa$, all of which were tested (data not shown): completely unconstrained (each center point as an individual parameter), uniformly distributed center points, center points distributed as roots of polynomials (e.g., shifted Chebyshev nodes – roots (also referred to as zeros) of Chebyshev polynomials, shifted to operate over the $[t_1,\ t_{N_t}]$ interval), and a version of the strategy above where we have several "focus" points (meaning we have several sin point "elements" or segments in the domain, i.e., we have multiple $\kappa$ fitted, where the number of $\kappa$ fitted is more than 1 but less than the total number of basis functions $N_f$). In other words, while using equation (15) produces 2 segments with a distribution of the center points, either side of $\kappa$ with the $p_1$ and $p_2$ coordinates, it is possible to have more segments. For example, three segments can be defined by two points shifted by $\kappa$.

#### ***Parameter and process variables used for the in-silico data generation***

The in-silico data were generated by simulating a culture fed under conditions analogous to the standard operations employed in Sartorius for CHO fed-batch processes. The culture is supplemented with three distinct feeds: FMA, FMB, and FMG. FMA consists of a mixture of amino acids and glucose, while FMB contains specific amino acids that are insoluble when combined with FMA. FMG is a concentrated glucose solution. The timing and volume of feed additions to the culture are detailed in Table S1. The glucose concentration in FMA is 75 g/L, and in FMG, it is 425 g/L. The culture is inoculated with 0.3 x $10^6$ cells/mL, accounting for a 1:10 ratio of dead cells, in a culture medium containing 6 g/L. Table S2 provides the parameter values utilized in the ODE model.

***Table S1**. Time and volume of feeding additions used to generate the in silico dataset*

| **FMA** | | **FMB** | | **FMG** | |
|---|---|---|---|---|---|
| **Time [days]** | **Volume [L]** | **Time [days]** | **Volume [L]** | **Time [days]** | **Volume [L]** |
| 3 | 0.007 | 3.01 | 0.0007 | | |
| 4 | 0.007 | 4.01 | 0.0007 | | |
| 5 | 0.0067 | 5.01 | 0.00067 | 5.1 | 0.0001 |
| 6 | 0.0065 | 6.01 | 0.00065 | 6.1 | 0.0001 |
| 7 | 0.0063 | 7.01 | 0.00063 | 7.1 | 0.00012 |
| 8 | 0.006 | 8.01 | 0.0006 | 8.05 | 0.0001 |
| 9 | 0.0058 | 9.01 | 0.00058 | 9.05 | 0.0008 |
| 10 | 0.0056 | 10.01 | 0.00056 | 10.05 | 0.0004 |
| 11 | 0.0054 | 11.01 | 0.00054 | 11.05 | 0.00015 |
| 12 | 0.0052 | 12.01 | 0.00052 | 12.05 | 0.0002 |
| 13 | 0.005 | 13.01 | 0.0005 | 13.05 | 0.0001 |
| 14 | 0.0048 | 14.01 | 0.00048 | | |

***Table S2**. ODE model parameters value*

| **Parameter** | **Description** | **Value** |
|---|---|---|
| $k_d$ | Death rate constant | 0.6 [day$^{-1}$] |
| $k_l$ | Lysis rate constant | 6 [day$^{-1}$] |
| $\mu_{G,max}$ | Maximum glucose consumption rate | 0.5 [$10^{-3}$ g ($10^6$ cells)$^{-1}$ day$^{-1}$] |
| $K_I$ | Half-saturation constant for inhibition by biomaterial | 10 [g.L$^{-1}$] |
| $m_G$ | Maintenance glucose uptake rate | 0.1[$10^{-3}$ g ($10^6$ cells)$^{-1}$ day$^{-1}$] |
| $K_G$ | Half-saturation constant for glucose uptake | 1[g.L$^{-1}$] |
| $\mu_{O,max}$ | Maximum oxidative capacity | 0.4 [$10^{-3}$ g ($10^6$ cells)$^{-1}$ day$^{-1}$] |
| $Y_L$ | Pseudo-stoichiometric coefficient for lactate formation | 6 [g.g$^{-1}$] |
| $K_L$ | Half-saturation constant for lactate uptake | 0.6 [g/L] |
| $Y_P$ | Pseudo-stoichiometric coefficient for protein product synthesis | 28 [$10^{-3}$ mg ($10^6$ cells)$^{-1}$ day$^{-1}$] |
| $Y_{v1}$ | Growth yield for oxidative metabolism mode | 4 [$10^3$ g$^{-1}$ ($10^6$ cells)] |
| $Y_{v2}$ | Growth yield for overflow metabolism mode | 0.3 [$10^3$ g$^{-1}$ ($10^6$ cells)] |
| $Y_{v3}$ | Growth yield for the lactate consumption mode | 1 [$10^3$ g$^{-1}$ ($10^6$ cells)] |

### Supplementary Figures

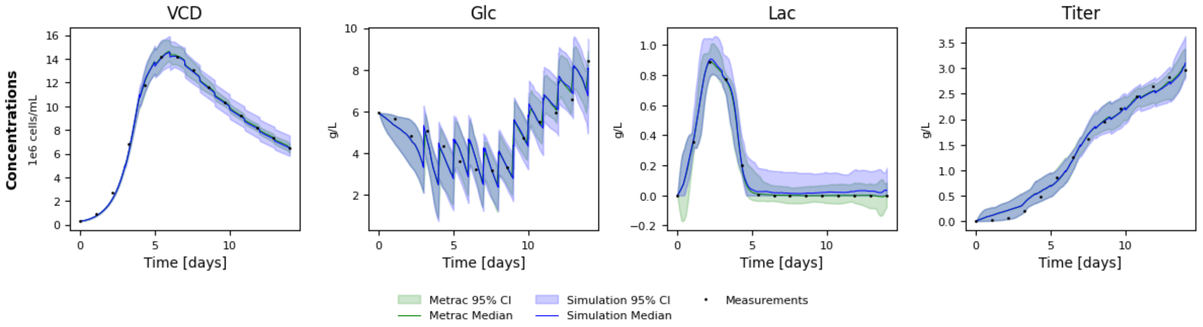


**Figure S1**. Comparison between MetRaC posterior distribution back-propagation and MetRaC-inferred rates into the ODE system (Eq. 45). Black dots denote simulated measurement data; green and blue shaded areas represent the 95% confidence intervals (CIs) of the MetRaC rate estimates and the resulting concentration simulations, respectively. Solid lines represent the corresponding medians.

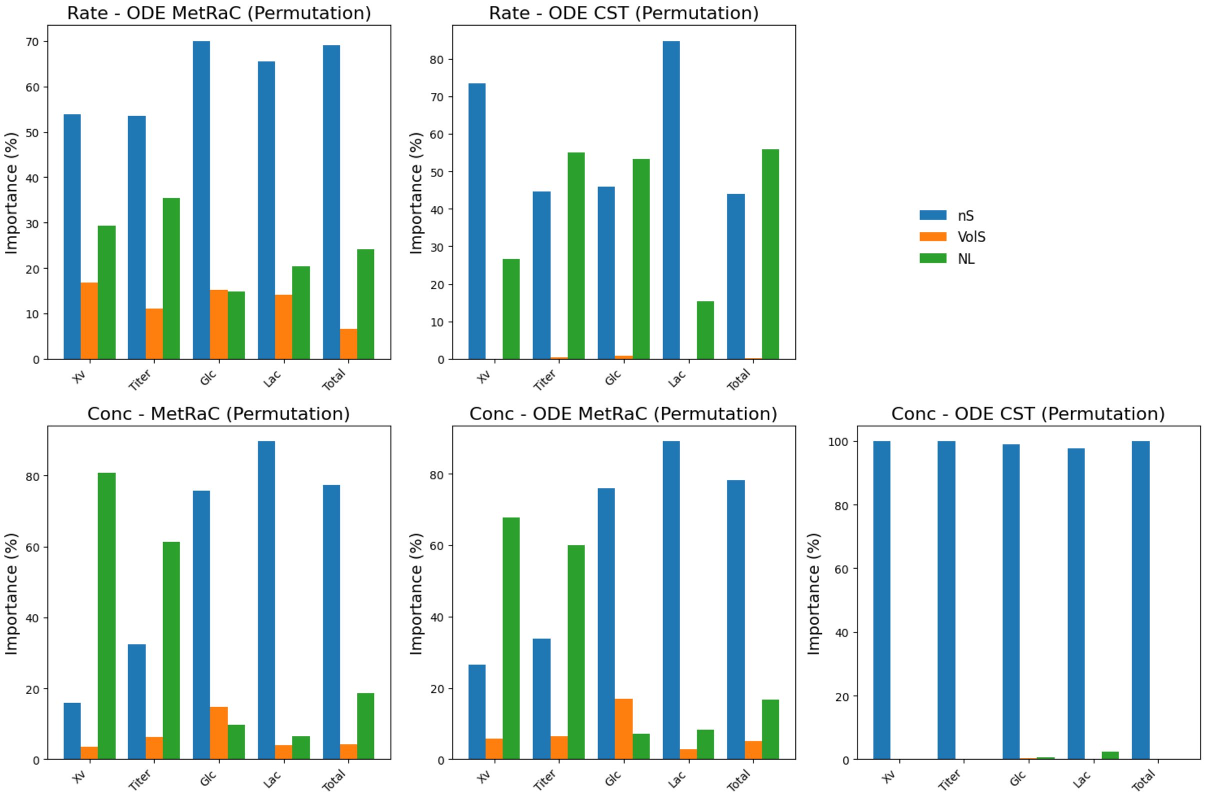


**Figure S2**. Relative importance of experimental design parameters on RMSE for rate estimation and concentration reconstruction. Importance was quantified using a Random Forest regression model (permutation-based approach) and expressed as the percentage of variance explained for each factor: sampling frequency (nS), sampling volume (VolS), and noise level (NL). Panels show results for: rate estimation using MetRaC (top left) and constant rate (top middle), concentration reconstruction via MetRaC backpropagation (bottom left), ODE simulation with MetRaC rates (bottom middle), and ODE simulation with constant rates (bottom right).
